# Phylogenetic relationships of *ATM* polymorphic loci haplotypes in Eurasian and African populations

**DOI:** 10.1101/2025.05.10.652950

**Authors:** Marina V. Olkova, Sergey M. Koshel, Georgiy Yu Ponomarev, Andrei A. Alimov

## Abstract

The *ATM* gene encodes a key kinase involved in DNA repair and cell cycle control, and mutations in it lead to the development of ataxia-telangiectasia and increase susceptibility to cancer. In this context, the study of the distribution patterns of *ATM* haplotypes in metapopulations living in Eurasia and Africa using population genetic and genogeographic methods appears to be a socially important scientific endeavor.

Polymorphic markers of the *ATM* gene were studied in 3876 individuals from 46 metapopulations, *ATM* gene haplotypes and genotypes based on *ATM* haplotypes were determined, and population frequencies of haplotypes and genotypes were calculated. An analysis of phylogenetic relationships of haplotypes, the Tajima test to assess population changes, and mapping of population frequencies of *ATM* gene haplotypes and genotypes were performed. The study showed the presence of a linkage block of 28 polymorphic markers of the *ATM* gene in the Eurasian and African populations. Two dominant *ATM* haplotypes in Eurasia, which are the result of–independent evolution of two branches from the ancestral *ATM* haplotype, are formed by mutually exclusive alleles and correspond to the “yin-yang” concept. The greatest diversity of *ATM* haplotypes is observed on the African continent. It is suggested that the process of haplotype formation occurred on the African continent, with subsequent migration events to the territory of Eurasia.

## INTRODUCTION

The human *ATM* gene (ataxia-telangiectasia mutated), which encodes a serine-threonine protein kinase, is located on chromosome 11, locus 11q22-23. Activation of the gene’s protein product occurs as a result of double-stranded DNA breaks, leading to the initiation of signaling pathways and the phosphorylation of numerous substrates (several hundred have been identified to date). The final result of cascade reactions is cell cycle arrest and DNA repair. In addition to double-stranded DNA breaks, activation of the *ATM* protein can be caused by oxidative stress, deficiency of a number of metabolites, hypoxia, and some other factors [1]. *ATM* is also involved in the activation of the spindle checkpoint during mitosis. Loss of *ATM* function results in a defective spindle checkpoint and increases the likelihood of aneuploidy [2].

Mutations in the *ATM* gene are associated with the development of autosomal recessive Louis-Barr syndrome (ataxia-telangiectasia) and with the risk of developing cancer, which is one of the reasons for the research interest in studying the distribution of frequencies of polymorphic variants of this gene in populations [3–6]. One of the most important stages of genetic data research is the linkage disequilibrium (LD) analysis, which allows to identify new clinically significant variants of the *ATM* gene associated with changes in the function of the gene’s protein product and disease severity. As a result of previous scientific studies, blocks of LD in the *ATM* gene have been identified and described in a region of ∼150 kb, covering the entire length of the gene [7–9]. In particular, Tornston et al. identified seven haplotypes of the *ATM* gene, two of which were present in 82% of the 93 DNA samples examined from representatives of seven human populations living on all continents. Two of the seven identified haplotypes were found only in African populations with a frequency of 33%. The presence of blocks of LD of varying length was also demonstrated for several other autosomal regions of the human genome. In particular, Bonnen’s article assessed the LD of non-coding polymorphic markers for 5 genes (TP53, RAG51, BRCA1, BRCA2 and *ATM*) associated with the cancer development. It was shown that for all genes, since the study included markers evenly distributed along the gene, there was no tendency for the LD value to decrease with increasing distance. The authors concluded that the LD value can vary widely regardless of the physical distribution of the markers along the length of the DNA sequence of the gene. The observed phenomenon testified to the existence of polymorphic loci in the genome, linked in blocks, with the degree of linkage independent of distance. A similar conclusion was reached by Gabriel et al., who reported the presence of SNP blocks with a small number of common haplotypes, evenly distributed across the human genome. The study was conducted using DNA samples from 275 representatives of the American population of African, European and Asian origin. A link was found between the origins of the populations and specific haplotypes. Haplotypes with mutually exclusive alleles of polymorphic markers, called “yin” and “yang”, were also identified [10–12].

Our study aims to perform a population analysis of combined into haplotypes SNP variants of the *ATM* gene, associated with oncological pathology. In addition, a genogeographic comparison of the North Eurasian gene pool of *ATM* haplotypes with the gene pools of the peoples of Eurasia and Africa was conducted. Phylogenetic analysis based on all identified *ATM* haplotypes was performed to better understand the structural changes accumulated in the gene during the evolutionary process.

## MATERIALS AND METHODS

### Ethical approvals

This study was conducted under the condition that written informed consent was obtained from all subjects. This consent was approved by the Ethics Committee of the Research Center for Medical Genetics.

### Collection of DNA genotype data and populations

1975 DNA samples from individuals representing 27 populations living in Russia and adjacent countries were provided by the Northern Eurasia Biobank. Population sample sizes ranged from 41 to 129 individuals. Individual allelic variants of 269 polymorphic markers of the *ATM* gene were extracted from a genome-wide genotyping dataset using the Infinium Omni5Exome BeadChip Kit (Illumina).

Genotype information for 1901 individuals from 19 populations in Eurasia and Africa, with population sample sizes ranging from 85 to 113 individuals, was obtained from the 1000 Genomes Project Phase 3 data (Ensembl).

The general list of populations, their abbreviated names and the size of population samples are presented in Table 1.

**Table 1.**
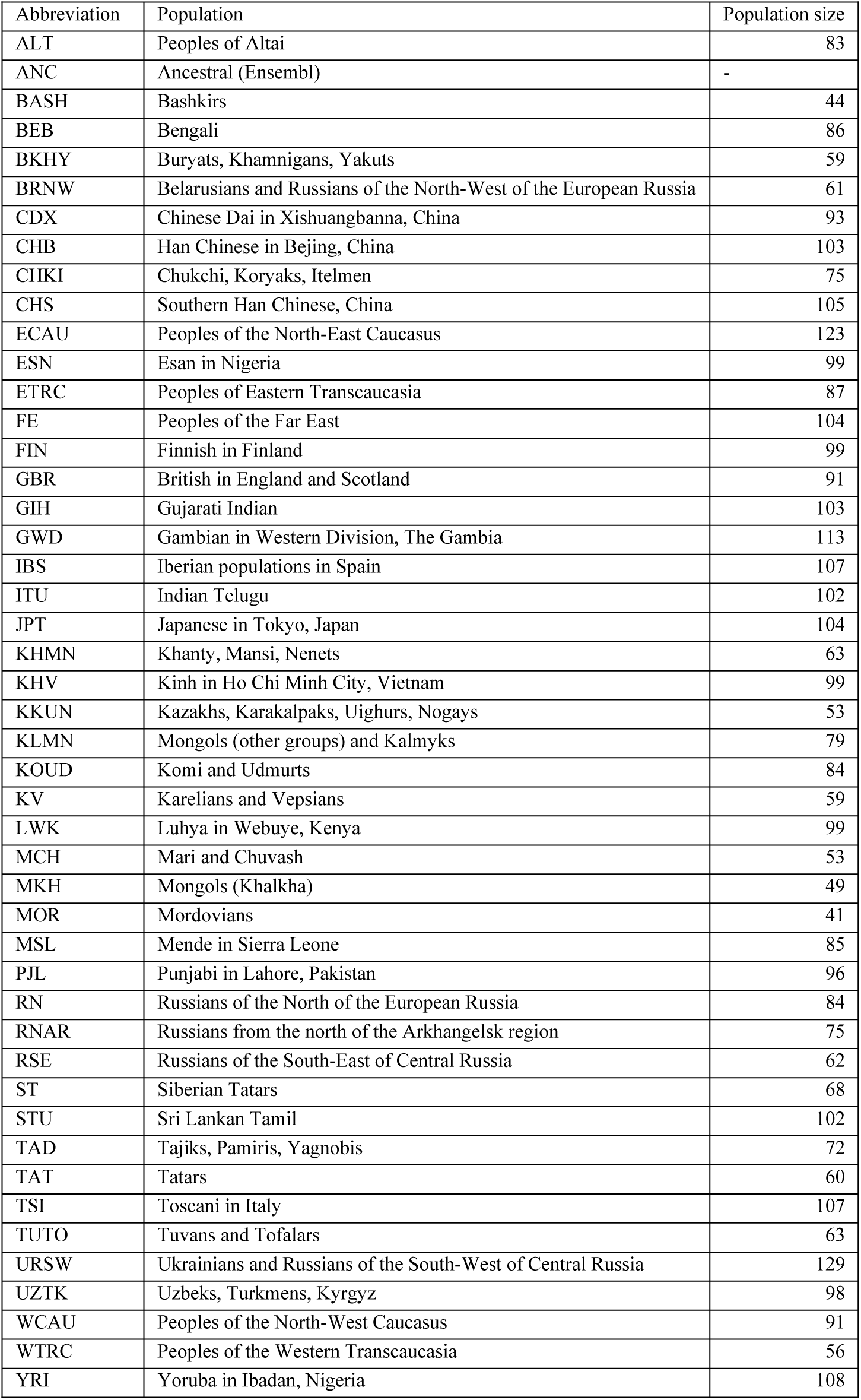
- Studied populations.

| Abbreviation | Population | Population size |
| --- | --- | --- |
| ALT | Peoples of Altai | 83 |
| ANC | Ancestral (Ensembl) | - |
| BASH | Bashkirs | 44 |
| BEB | Bengali | 86 |
| BKHY | Buryats, Khamnigans, Yakuts | 59 |
| BRNW | Belarusians and Russians of the North-West of the European Russia | 61 |
| CDX | Chinese Dai in Xishuangbanna, China | 93 |
| CHB | Han Chinese in Beijing, China | 103 |
| CHKI | Chukchi, Koryaks, Itelmen | 75 |
| CHS | Southern Han Chinese, China | 105 |
| ECAU | Peoples of the North-East Caucasus | 123 |
| ESN | Esan in Nigeria | 99 |
| ETRC | Peoples of Eastern Transcaucasia | 87 |
| FE | Peoples of the Far East | 104 |
| FIN | Finnish in Finland | 99 |
| GBR | British in England and Scotland | 91 |
| GIH | Gujarati Indian | 103 |
| GWD | Gambian in Western Division, The Gambia | 113 |
| IBS | Iberian populations in Spain | 107 |
| ITU | Indian Telugu | 102 |
| JPT | Japanese in Tokyo, Japan | 104 |
| KHMN | Khanty, Mansi, Nenets | 63 |
| KHV | Kinh in Ho Chi Minh City, Vietnam | 99 |
| KKUN | Kazakhs, Karakalpaks, Uighurs, Nogays | 53 |
| KLMN | Mongols (other groups) and Kalmyks | 79 |
| KOUD | Komi and Udmurts | 84 |
| KV | Karelians and Vepsians | 59 |
| LWK | Luhya in Webuye, Kenya | 99 |
| MCH | Mari and Chuvash | 53 |
| MKH | Mongols (Khalkha) | 49 |
| MOR | Mordovians | 41 |
| MSL | Mende in Sierra Leone | 85 |
| PJL | Punjabi in Lahore, Pakistan | 96 |
| RN | Russians of the North of the European Russia | 84 |
| RNAR | Russians from the north of the Arkhangelsk region | 75 |
| RSE | Russians of the South-East of Central Russia | 62 |
| ST | Siberian Tatars | 68 |
| STU | Sri Lankan Tamil | 102 |
| TAD | Tajiks, Pamiris, Yagnobis | 72 |
| TAT | Tatars | 60 |
| TSI | Toscani in Italy | 107 |
| TUTO | Tuvans and Tofalars | 63 |
| URSW | Ukrainians and Russians of the South-West of Central Russia | 129 |
| UZTK | Uzbeks, Turkmen, Kyrgyz | 98 |
| WCAU | Peoples of the North-West Caucasus | 91 |
| WTRC | Peoples of the Western Transcaucasia | 56 |
| YRI | Yoruba in Ibadan, Nigeria | 108 |

The “ancestral” haplotype was determined based on the “ancestral” alleles of the polymorphic markers specified in the Ensembl project, identified based on a three-stage multiple alignment of EPO (Enredo, Pecan, Ortheus).

### Statistical data processing

Microsoft Excel (Microsoft Corporation) and RStudio 2024.12.1+563 - R version 4.4.2 (RStudio, PBC) were used for statistical data processing.

### DNA sample genotyping and polymorphic ATM loci quality assessment

Polymorphic markers and samples with genotype quality below 95% were excluded from the study (using the *snpStats* R-package).

### Isolation of haplotypes. Calculation of population frequencies of *ATM* haplotypes and genotypes

The isolation of haplotypes and the calculation of their population frequencies were performed in the Haploview 4.1 program (Broad Institute).

The “confidence interval method” (Gabriel) was used to isolate blocks of polymorphic variants forming a haplotype. A haplotype block was defined if 95% of SNP pairs within it had D > 0.8. Calculation of population frequencies of genotypes based on *ATM* haplotypes was performed using the *plyr* R-package.

### Phylogenetic analysis

Phylogenetic networks of *ATM* haplotype representation in populations were constructed using the *pegas* and *geneHapR* R-packages according to the “parsimony principle”. The preparatory files were filled with data on population frequencies of haplotypes, multiplied by 1000.

### Calculation of Tajima test for populations and individual “yin” and “yang” branches

The Tajima test was calculated based on the population haplotype frequency data obtained in Haploview 4.1 software. The population number of haplotypes of each type was obtained by multiplying the haplotype population frequency by 1000. The Tajima test D-value and p-value were calculated using the *pegas* R-package.

### Cartographic analysis

Cartographic visualization of the population frequencies of haplotypes and genotypes of the *ATM* gene was performed using the original GeneGeo cartographic package [13]. Frequency calculations for the entire mapped area were performed using the weighted average interpolation method with a weighting function inversely proportional to the cube of the distance and an influence radius of 4000 km. To avoid the influence of map projection distortions calculations, all distances in the interpolation process were calculated on a sphere. The Winkel triple projection with the central meridian 60° E was used to create the maps.

## RESULTS

### Determination of *ATM* gene haplotypes and their population frequencies

The genotype data of the combined population of Northern Eurasia allowed the identification of a LD block of 28 linked polymorphic loci using Haploview 4.1 software – rs189037, rs228590, rs623860, rs600931, rs228592, rs672655, rs627418, rs664677, rs609261, rs645485, rs619972, rs599558, rs595747, rs609655, rs227061, rs227062, rs227064, rs227068, rs227070, rs227074, rs227075, rs425538, rs419716, rs227040, rs664143, rs227094, rs227092 and rs4585, located over 145,795 bp from the 5′-untranslated region to the 3′-untranslated region of the *ATM* gene (Figure S1).

The value of the square correlation coefficient (r^2^) between all pairs of loci in the LD block exceeded 0.8. Of 268,435,456 possible haplotypes for 28 polymorphic loci of the LD block in the combined population of peoples living in the territory of Northern Eurasia, 4 haplotypes were found, and 2 of them were predominant: G-A-C-C-C-A-T-C-T-A-C-C-C-G-A-G-T-A-T-A-T-C-A-T-A-C-G-G and A-G-T-T-A-G-C-T-C-G-T-T-T-A-G-A-C-G-G-G-C-A-C-C-G-A-T-T with a frequency of 39.2% and 54.5%, respectively. The alleles in the two most frequent haplotypes were mutually exclusive, which allows us to classify them as “yin”/“yang” haplotypes. The remaining two detected haplotypes were much less frequent in the population (with a frequency of about 2%) and were the result of recombination between the “yin” and “yang” haplotypes.

Similarly, haplotypes were determined, and their population frequencies were calculated for 28 polymorphic markers of the *ATM* haplotype block in each of the 27 populations of Northern Eurasia and in 19 populations of Eurasia and Africa using data from the 1000 Genomes Project Phase 3 (Ensembl). From the total set of haplotypes detected in all the above Eurasian and African populations, rare (with a population frequency of up to 5%) found in single populations, as well as haplotypes that most likely arose as a result of recombination between the “yin-yang” haplotypes, were excluded. As a result, a list of 11 haplotypes of the *ATM* gene (Figure 1) was determined, occurring in 46 populations of Eurasia and Africa. The “ancestral” haplotype (H0) of the human *ATM* gene was determined using the Ensembl genome browser, which indicates the “ancestral” alleles for each of the 28 linked loci of this gene.

**Figure 1.**
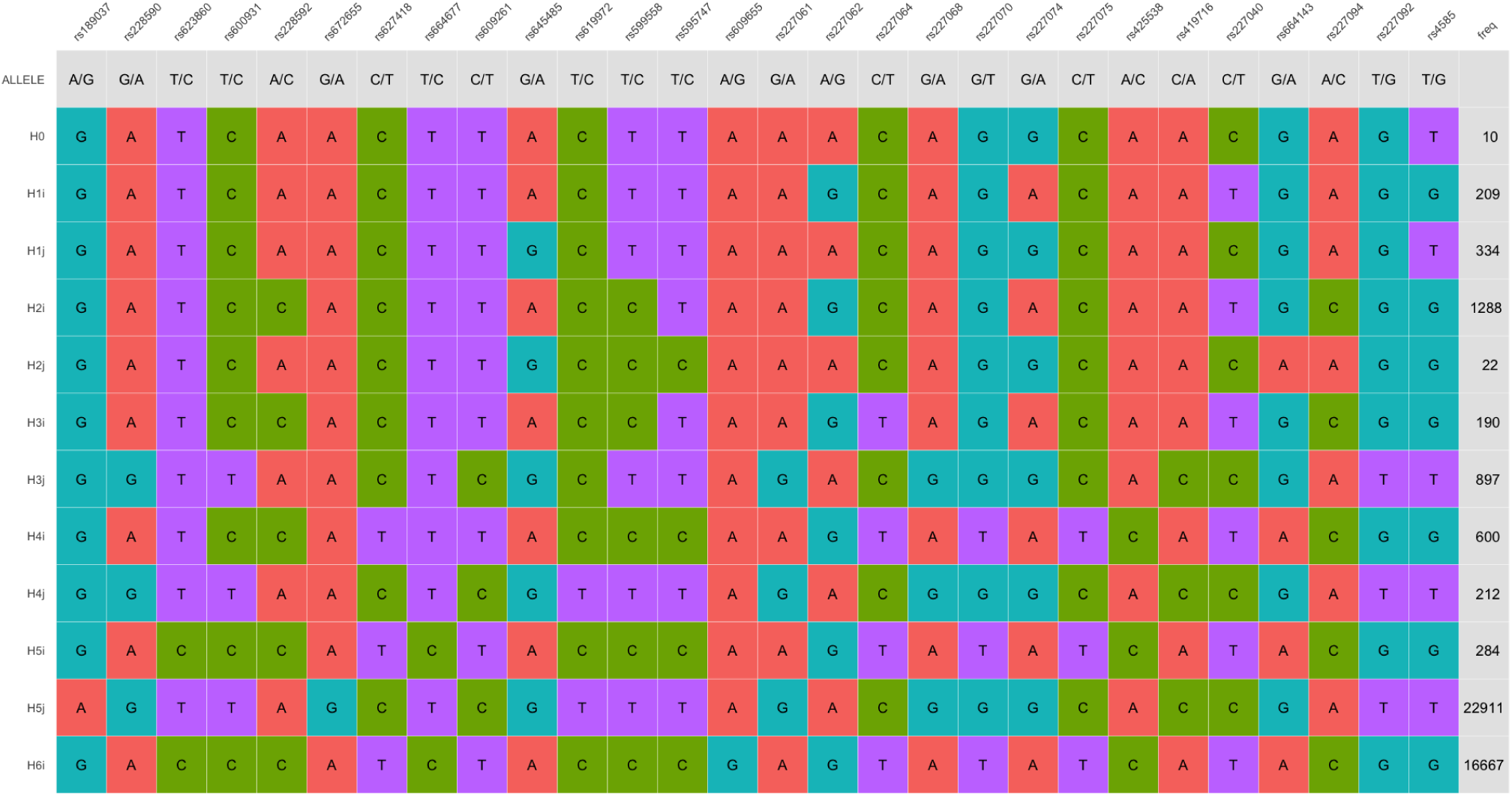
– *ATM* haplotypes.

### *ATM* haplotypes-based phylogenetic analysis

Based on the isolated haplotypes, a phylogenetic network was constructed reflecting the evolutionary variability of the *ATM* gene haplotype block in the human population and the frequency ratio of *ATM* haplotypes in modern peoples of Eurasia and Africa (Figure 2).

**Figure 2.**
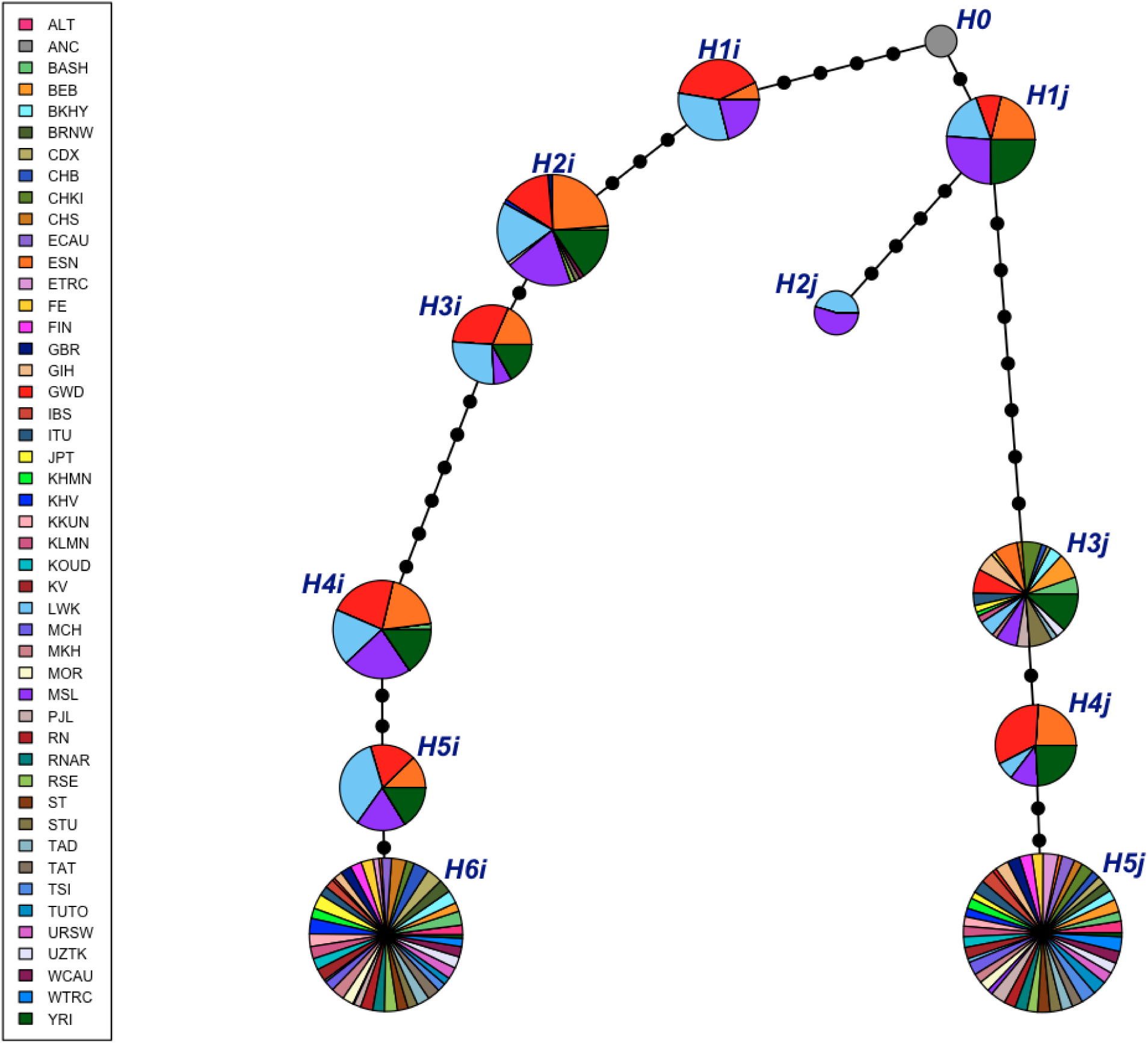
- Phylogenetic network of *ATM* haplotypes in Eurasian and African populations

The presence of two independently evolving phylogenetic branches, which we called I and J branches, was discovered, leading from the “ancestral” haplotype to the most common in Eurasia H6i and H5j (“yin” and “yang”) haplotypes (Figure 3). All haplotypes analyzed were found in African populations (Figure 4, S2a).

**Figure 3.**
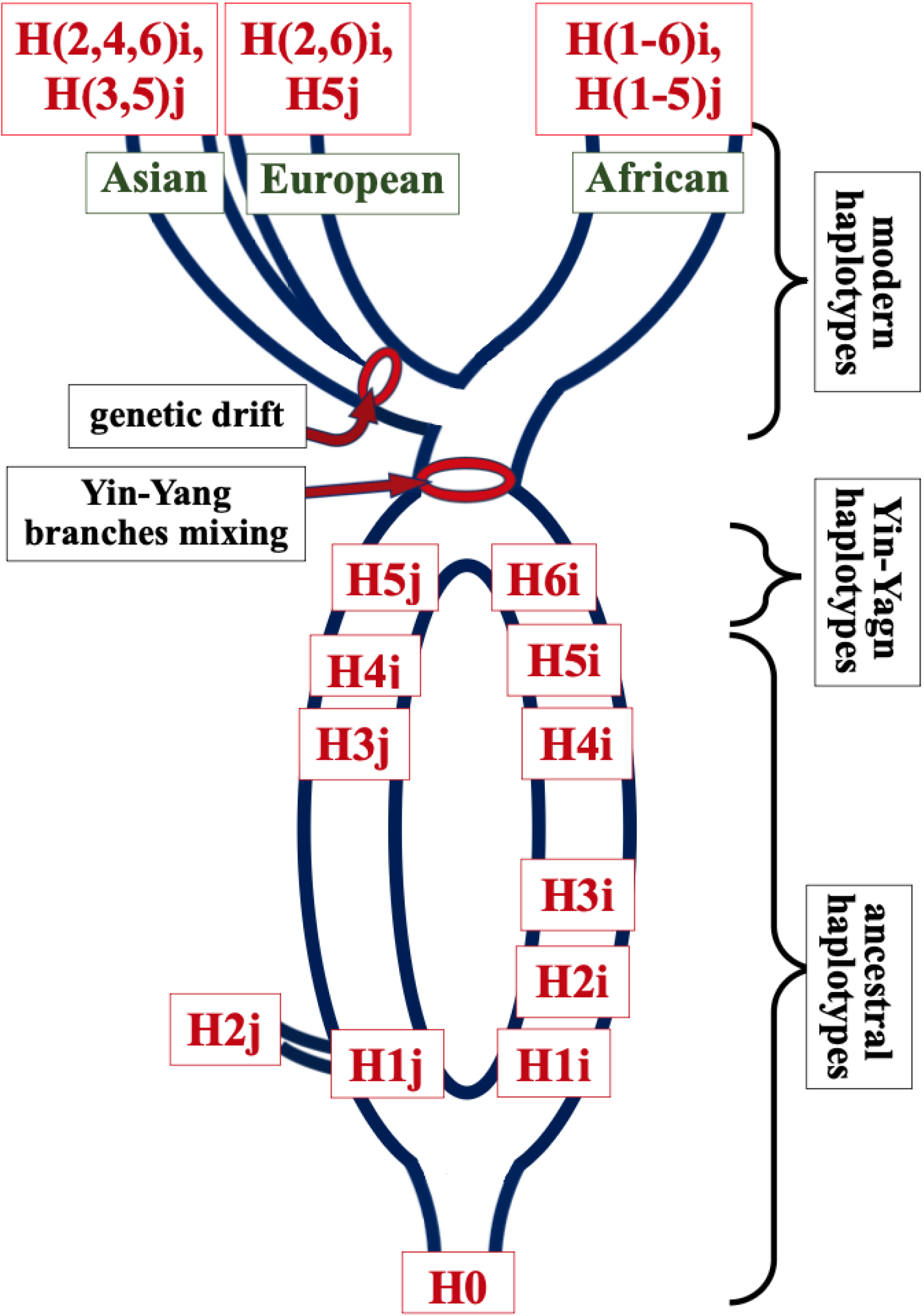
- Scheme of phylogenetic relationships of *ATM* haplotypes

**Figure 4.**
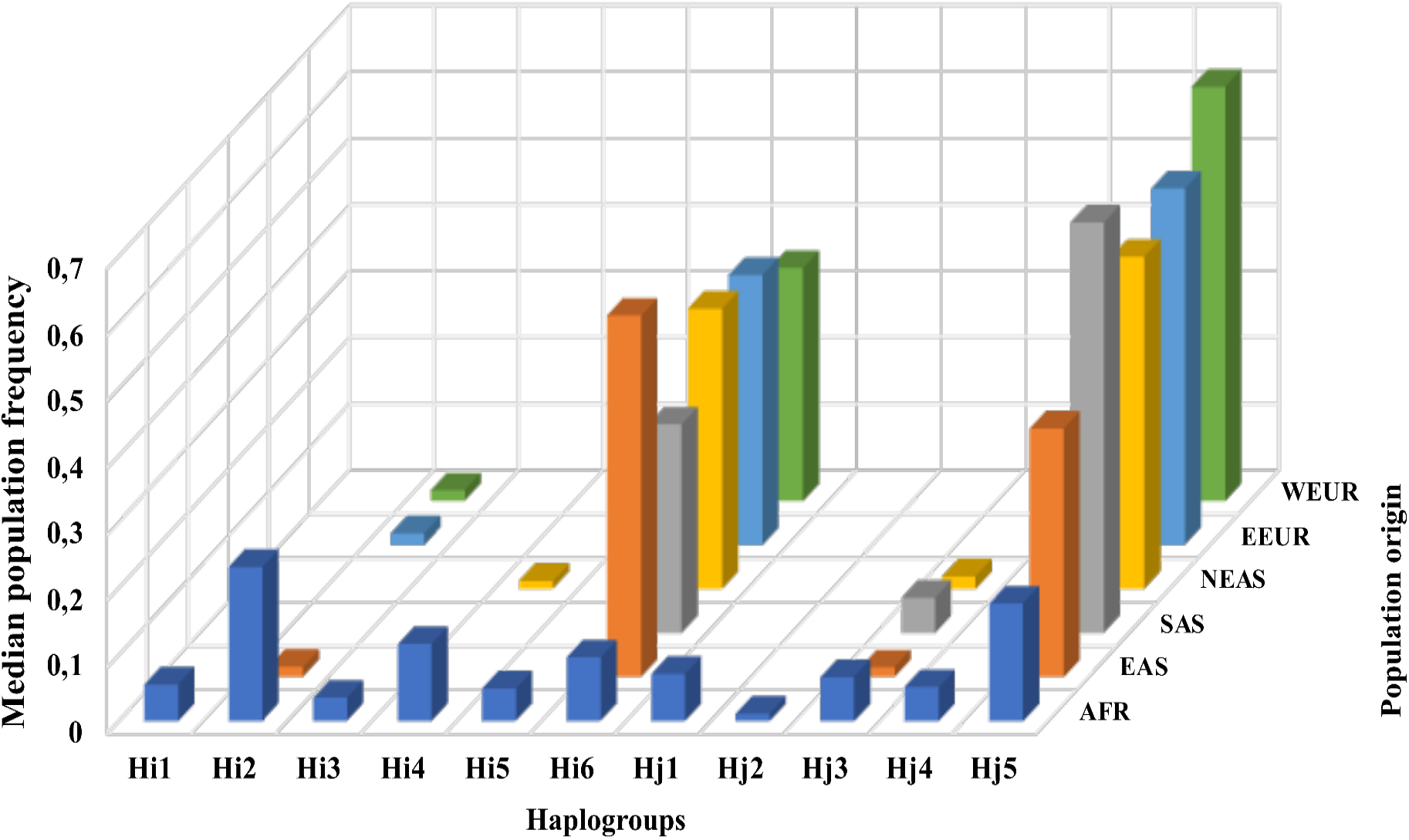
– Medians of population frequencies of *ATM* haplotypes in populations by origin: AFR – African, EAS – East Asian, EEUR – Eastern European, NEAS – Asian of Northern Eurasia, SAS – South Asian, WEUR – Western European

In addition to the main “yin/yang” haplotypes, the H2i and H4i (branch I), H3j (branch J) haplotypes were also present in the Eurasia area. Of these three haplotypes, only H2i was found in populations of European origin (Figure S2b), in contrast to populations of Asian origin (Figure S2c). All three haplotypes are rare (up to 2%) in Eurasian populations. Based on these data, it can be concluded that the formation of the ancestral haplotype and the subsequent divergent evolution of two isolated branches probably occurred on the territory of the African continent.

### Tajima test for populations and individual branches of “yin” and “yang” haplotypes

For all populations of Eurasia and Africa, both divided by origin and geographical location, and combined by these criteria, a positive value of the D parameter of the Tajima test (from 4.7 to 9.3) with a p value << 0.05 was observed, which may be due to a long-term subdivision of the population into several subgroups with limited gene flow between them, balancing selection, a sharp reduction in population size, or other genetic and demographic processes.

For each of these populations, a significant deviation from the neutral model of evolution was observed. At the same time, when considering each of the two evolutionary branches leading from the “ancestral” haplotype to the H6i and H5j haplotypes separately, this test showed a statistically insignificant result for the D parameter and p-value (p >> 0.05), which indicating that both divergence paths (I and J) of the “ancestral” *ATM* haplotype conform to the neutral model of evolution (Table S1).

### Geographical analysis of population frequencies of *ATM* haplotypes and genotypes

The frequencies of *ATM* haplotypes, as well as their homozygous and heterozygous combinations, are presented in Tables 2-3 and on maps (Fig. S3-S6).

**Table 2.** - Population frequencies of *ATM* haplotypes. See Table 1 for population abbreviations

| Population | H1i | H1j | H2i | H2j | H3i | H3j | H4i | H4j | H5i | H5j | H6i |
| --- | --- | --- | --- | --- | --- | --- | --- | --- | --- | --- | --- |
| ALT |  |  |  |  |  |  |  |  |  | 0.556 | 0.344 |
| BASH |  |  |  |  |  | 0.048 | 0.012 |  |  | 0.440 | 0.488 |
| BEB |  |  |  |  |  | 0.070 |  |  |  | 0.587 | 0.314 |
| BKHY |  |  |  |  |  | 0.036 |  |  |  | 0.464 | 0.455 |
| BRNW |  |  |  |  |  |  |  |  |  | 0.458 | 0.467 |
| CDX |  |  | 0.016 |  |  | 0.011 |  |  |  | 0.360 | 0.559 |
| CHB |  |  |  |  |  | 0.015 |  |  |  | 0.374 | 0.544 |
| CHKI |  |  |  |  |  | 0.055 |  |  |  | 0.591 | 0.280 |
| CHS |  |  |  |  |  | 0.014 |  |  |  | 0.390 | 0.524 |
| ECAU |  |  |  |  |  |  |  |  |  | 0.604 | 0.346 |
| ESN | 0.015 | 0.071 | 0.308 |  | 0.035 | 0.066 | 0.116 | 0.051 | 0.035 | 0.177 | 0.096 |
| ETRC |  |  |  |  |  |  |  |  |  | 0.715 | 0.209 |
| FE |  |  |  |  |  | 0.011 |  |  |  | 0.517 | 0.422 |
| FIN |  |  |  |  |  |  |  |  |  | 0.581 | 0.399 |
| GBR |  |  | 0.016 |  |  |  |  |  |  | 0.588 | 0.379 |
| GIH |  |  |  |  |  | 0.053 |  |  |  | 0.631 | 0.306 |
| GWD | 0.084 | 0.031 | 0.186 |  | 0.058 | 0.067 | 0.133 | 0.071 | 0.049 | 0.177 | 0.097 |
| IBS |  |  |  |  |  |  |  |  |  | 0.659 | 0.322 |
| ITU |  |  |  |  |  | 0.034 |  |  |  | 0.618 | 0.333 |
| JPT |  |  |  |  |  | 0.019 |  |  |  | 0.322 | 0.514 |
| KHMN |  |  |  |  |  | 0.012 |  |  |  | 0.492 | 0.359 |
| KHV |  |  | 0.015 |  |  |  |  |  |  | 0.399 | 0.545 |
| KKUN |  |  |  |  |  |  |  |  |  | 0.442 | 0.450 |
| KLMN |  |  |  |  |  | 0.019 |  |  |  | 0.494 | 0.430 |
| KOUD |  |  |  |  |  |  |  |  |  | 0.499 | 0.402 |
| KV |  |  |  |  |  |  |  |  |  | 0.533 | 0.440 |
| LWK | 0.066 | 0.061 | 0.232 | 0.010 | 0.051 | 0.045 | 0.111 | 0.015 | 0.101 | 0.182 | 0.076 |
| MCH |  |  |  |  |  |  |  |  |  | 0.616 | 0.353 |
| MKH |  |  |  |  |  | 0.014 |  |  |  | 0.443 | 0.471 |
| MOR |  |  | 0.012 |  |  |  |  |  |  | 0.499 | 0.390 |
| MSL | 0.044 | 0.088 | 0.250 | 0.012 | 0.014 | 0.059 | 0.135 | 0.024 | 0.053 | 0.241 | 0.041 |
| PJL |  |  |  |  |  | 0.036 |  |  |  | 0.703 | 0.250 |
| RN |  |  |  |  |  |  |  |  |  | 0.537 | 0.431 |
| RNAR |  |  |  |  |  |  |  |  |  | 0.568 | 0.412 |
| RSE |  |  | 0.018 |  |  |  |  |  |  | 0.500 | 0.456 |
| ST |  |  |  |  |  |  |  |  |  | 0.571 | 0.375 |
| STU |  |  |  |  |  | 0.064 |  |  |  | 0.554 | 0.353 |
| TAD |  |  |  |  |  | 0.017 |  |  |  | 0.517 | 0.441 |
| TAT |  |  | 0.019 |  |  |  |  |  |  | 0.497 | 0.442 |
| TSI |  |  |  |  |  |  |  |  |  | 0.692 | 0.299 |
| TUTO |  |  |  |  |  |  |  |  |  | 0.659 | 0.254 |
| URSW |  |  |  |  |  |  |  |  |  | 0.536 | 0.411 |
| UZTK |  |  |  |  |  | 0.026 |  |  |  | 0.500 | 0.408 |
| WCAU |  |  | 0.017 |  |  |  |  |  |  | 0.565 | 0.365 |
| WTRC |  |  |  |  |  |  |  |  |  | 0.687 | 0.295 |
| YRI |  | 0.083 | 0.199 |  | 0.032 | 0.106 | 0.093 | 0.051 | 0.046 | 0.176 | 0.120 |

**Table 3.** - Population genotype frequencies based on *ATM* haplotypes. See Table 1 for population abbreviations

| Population | Population frequency of homozygous genotype |  |  |  |  |  |  |  |  | Population frequency of heterozygous genotype |  |  |  |  |  |
| --- | --- | --- | --- | --- | --- | --- | --- | --- | --- | --- | --- | --- | --- | --- | --- |
|  | H6i/<br>H6i | H5j/<br>H5j | H4i/<br>H4i | H3i/<br>H3i | H2i/<br>H2i | H1j/<br>H1j | H1i/<br>H1i | H5i/<br>H5i | H4j/<br>H4j | H6i/<br>H5j | H2i/<br>H6i | H2i/<br>H5j | H3j/<br>H6i | H3j/<br>H5j | H4i/<br>H2i |
| ALT | 0.088 | 0.238 |  |  |  |  |  |  |  | 0.400 |  |  | 0.013 |  |  |
| BASH | 0.238 | 0.143 |  |  |  |  |  |  |  | 0.357 |  |  | 0.071 | 0.024 |  |
| BEB | 0.116 | 0.337 |  |  |  |  |  |  |  | 0.349 |  |  | 0.035 | 0.105 |  |
| BKHY | 0.232 | 0.161 |  |  |  |  |  |  |  | 0.286 | 0.018 |  | 0.036 | 0.018 |  |
| BRNW | 0.217 | 0.250 |  |  |  |  |  |  |  | 0.383 |  |  |  |  |  |
| CDX | 0.333 | 0.151 |  |  |  |  |  |  |  | 0.355 | 0.022 | 0.011 | 0.011 | 0.011 | 0.022 |
| CHB | 0.340 | 0.175 |  |  |  |  |  |  |  | 0.330 |  | 0.010 | 0.019 | 0.010 |  |
| CHKI | 0.145 | 0.345 |  |  |  |  |  |  |  | 0.164 |  |  | 0.036 | 0.055 |  |
| CHS | 0.267 | 0.114 |  |  |  |  |  |  |  | 0.448 | 0.010 | 0.010 | 0.019 | 0.010 | 0.010 |
| ECAU | 0.123 | 0.298 |  |  |  |  |  |  |  | 0.360 |  | 0.009 |  |  |  |
| ESN | 0.020 | 0.040 | 0.020 | 0.010 | 0.101 |  |  |  |  | 0.010 | 0.051 | 0.071 | 0.010 | 0.020 | 0.051 |
| ETRC | 0.012 | 0.420 |  |  |  |  |  |  |  | 0.272 |  |  |  |  |  |
| FE | 0.178 | 0.211 |  |  |  |  |  |  |  | 0.378 |  |  | 0.011 | 0.011 |  |
| FIN | 0.202 | 0.364 |  |  |  |  |  |  |  | 0.394 |  | 0.010 |  |  |  |
| GBR | 0.121 | 0.341 |  |  |  |  |  |  |  | 0.473 | 0.022 | 0.011 |  |  | 0.022 |
| GIH | 0.097 | 0.417 |  |  |  |  |  |  |  | 0.359 |  | 0.010 | 0.058 | 0.049 |  |
| GWD | 0.009 | 0.035 | 0.044 | 0.009 | 0.018 |  | 0.009 |  | 0.009 | 0.044 | 0.018 | 0.053 | 0.018 | 0.027 | 0.018 |
| IBS | 0.084 | 0.411 |  |  |  |  |  |  |  | 0.467 |  | 0.009 |  |  |  |
| ITU | 0.108 | 0.382 |  |  |  |  |  |  |  | 0.422 |  |  | 0.020 | 0.039 |  |
| JPT | 0.279 | 0.115 |  |  |  |  |  |  |  | 0.279 |  | 0.019 | 0.029 | 0.010 |  |
| KHMN | 0.097 | 0.210 |  |  |  |  |  |  |  | 0.339 |  |  | 0.016 |  |  |
| KHV | 0.263 | 0.101 |  |  |  |  |  |  |  | 0.525 | 0.010 | 0.020 | 0.010 |  | 0.010 |
| KKUN | 0.154 | 0.154 |  |  |  |  |  |  |  | 0.404 |  |  | 0.019 |  |  |
| KLMN | 0.203 | 0.215 |  |  |  |  |  |  |  | 0.342 |  |  | 0.013 | 0.025 |  |
| KOUD | 0.159 | 0.244 |  |  |  |  |  |  |  | 0.341 | 0.017 |  |  |  |  |
| KV | 0.224 | 0.328 |  |  |  |  |  |  |  | 0.362 |  |  |  |  |  |
| LWK |  | 0.030 | 0.020 |  | 0.061 |  | 0.010 | 0.010 |  | 0.030 | 0.040 | 0.051 |  | 0.020 | 0.040 |
| MCH | 0.137 | 0.353 |  |  |  |  |  |  |  | 0.294 |  |  | 0.020 |  |  |
| MKH | 0.200 | 0.244 |  |  |  |  |  |  |  | 0.356 |  |  |  | 0.022 |  |
| MOR | 0.098 | 0.244 |  |  |  |  |  |  |  | 0.317 | 0.024 |  |  |  |  |
| MSL |  | 0.059 | 0.024 |  | 0.082 | 0.035 |  |  |  | 0.024 | 0.047 | 0.071 |  | 0.047 | 0.047 |
| PJL | 0.083 | 0.510 |  |  |  |  |  |  |  | 0.313 |  |  | 0.010 | 0.063 |  |
| RN | 0.213 | 0.338 |  |  |  |  |  |  |  | 0.363 |  | 0.013 |  |  |  |
| RNAR | 0.135 | 0.297 |  |  |  |  |  |  |  | 0.514 |  |  |  |  |  |
| RSE | 0.211 | 0.228 |  |  |  |  |  |  |  | 0.421 | 0.018 | 0.018 |  | 0.018 |  |
| ST | 0.107 | 0.250 |  |  |  |  |  |  |  | 0.429 |  |  |  |  |  |
| STU | 0.147 | 0.363 |  |  |  |  |  |  |  | 0.314 |  |  | 0.078 | 0.039 |  |
| TAD | 0.136 | 0.220 |  |  |  |  |  |  |  | 0.424 |  |  | 0.034 |  |  |
| TAT | 0.154 | 0.173 |  |  |  |  |  |  |  | 0.481 |  | 0.038 |  |  |  |
| TSI | 0.121 | 0.514 |  |  |  |  |  |  |  | 0.346 | 0.009 | 0.009 |  |  | 0.009 |
| TUTO | 0.063 | 0.413 |  |  |  |  |  |  |  | 0.254 |  |  |  | 0.016 |  |
| URSW | 0.195 | 0.285 |  |  |  |  |  |  |  | 0.350 |  |  |  |  |  |
| UZTK | 0.143 | 0.235 |  |  |  |  |  |  |  | 0.398 | 0.010 |  | 0.020 | 0.031 |  |
| WCAU | 0.101 | 0.292 |  |  |  |  |  |  |  | 0.360 | 0.011 |  |  |  |  |
| WTRC | 0.071 | 0.446 |  |  |  |  |  |  |  | 0.411 |  |  |  | 0.018 |  |
| YRI |  | 0.009 | 0.009 |  | 0.019 |  |  |  |  | 0.065 | 0.046 | 0.046 | 0.046 | 0.037 | 0.046 |

### “Yin” and “yang” haplotypes

Cartographic analysis of the prevalence of population frequencies of haplotypes H6i and H5j, as well as homozygous carriers of these haplotypes showed similar results.

Haplotype H5j was in southern Eurasia with maxima in the southern of Western Europe (∼ 69%), in Transcaucasia (∼72%) and in the north-west of South Asia (∼70%). There was also a significant increase in the frequency of this haplotype in the populations of the indigenous populations of southern Siberia (with a maximum among Tuvans and Tofalars ∼66%) and Kamchatka (Chukchi, Koryaks, Itelmens ∼59%), with a significant decrease in East Asia to a frequency of ∼36%. The H5j haplotype is the second most common in Africa, although its population frequencies are noticeable lower than in Eurasia - from 18% in most African populations to 24% in the Mende people living on the coast of West Africa.

The H6i haplotype has two geographic maxima - on the east coast of Asia (∼56% in the Dai people in southwest China) and in eastern Europe (∼49% in the Bashkirs). Between the maxima, there is a marked decrease in population frequencies in central Siberia, extending to the northern coast of Eurasia and the south of the continent - along the northern border of the Alpide belt. The lowest population frequency of the H6i haplotype in Eurasia is in Eastern Transcaucasia (∼21%). The H6i haplotype is the fourth most common in African populations – from 4% in the Mende of Sierra Leone to 12% in the Yoruba of Nigeria.

In summary, it can be said that in the Eurasian area, apart from East Asia, the H5j haplotype predominates between the two mutually exclusive and most common haplotypes. In the populations of East Asia, the H6i haplotype predominates. In Africa, the population frequencies of the “yin” and “yang” haplotypes are significantly lower than in Eurasian populations. On average, the H5j haplotype is 2.8 times less common in African populations than in Eurasian populations, and the H6i haplotype is 4.6 times less common.

### Heterozygous combination of “yin” and “yang” haplotypes

The heterozygous genotype H6i/H5j turned out was one of the most common in Eurasian populations (from 16.4% to 52.5%). The exception was 9 out of 42 Eurasian populations (Table 5). The prevalence of homozygotes for the H5j haplotype was found in 2 populations of northwestern South Asia (41.7% in Gujaratis of India and 51% in Punjabis of Pakistan), in populations of Transcaucasia (42-44.6%), in the Italian population of southern Europe (51.4%), as well as in 3 metapopulations of Northern Eurasia - “Chukchi, Koryaks, Itelmens” (34.5%), “Mari, Chuvash” (35.3%) and “Tuvans, Tofalars” (41.3%). The homozygous genotype for the H6i haplotype turned out was the most common in only one population - the East Asian Han, where its frequency was ∼ 34%. In Africa, the frequency of H6i/H5j heterozygotes was an order of magnitude lower than in Eurasian (on average ∼3.5%) than in Eurasia (∼37%). Meanwhile, among the Yoruba population, the genotype H6i/H5j was the most frequent (6.5%), despite the complete absence of homozygotes for haplotype H6i and a very low population frequency of homozygotes for haplotype H5j (∼1%). This may be related to the manifestation of balancing selection or migration processes.

The peaks of the H6i/H5j genotype in Eurasia are located on the coasts: in the east of the Indochinese peninsula (∼52.5%), in the Spanish (∼46.7%) and British (∼47.3%) populations of Western Europe, and in the Arkhangelsk region (∼51.4%) of Northern Europe. There is also a significant increase in heterozygote frequencies in the Eastern European Tatar population (48.1%).

The Eurasian minimum of population frequencies of “yin-yang” heterozygotes is observed among the “Chukchi, Koryaks and Itelmens” (∼16.4%).

### Haplotype H3j

Haplotype H3j differs from H5j at 3 positions - rs189037, rs672655, rs619972. With a frequency of 1% to 7%, H3j is found in most Eurasian populations of Asian origin (except for the West Siberian Plain and Vietnam) but has not been found in European populations. Homozygotes for H3j have not been found, apparently because of the low frequencies of the haplotype.

In Africa, this haplotype ranks 5th in population frequency ranging - from 4.5% in the Luhya population of Kenya to 11% in the Yoruba of Nigeria.

### Geography of population frequencies of heterozygotes for the haplotype H3j in combination with H6i, H5j, H4i and H2i

The geographical distribution of the H3j haplotype in combination with H6i and H5j generally corresponds to that of the H3j haplotype in populations. The H3j/H6i genotypes are more common along the line running from southern Asia to northern Eurasia, with the highest frequencies among Sri Lankan Tamils (7.8%) and Bashkirs (7.1%). The H3j/H5j genotype is observed with the highest frequency (10.5%) in the Bengal population of South Asia.

The H3j/H4i and H3j/H2i genotypes are found only in Africa, the former with a frequency of 1-2% and the latter with a frequency of 2-6%.

### Haplotype H2i

The most common in Africa, H2i (ranging from 19% to 31%), was found at very low levels (∼2%) in some populations of East Asia (southern China and Vietnam) and Europe (UK, Western Caucasus, south-eastern central Russia, Mordovia and Tatarstan).

### Prevalence of heterozygous carriers of the haplotype H2i in combination with H6i, H5j and H4i

The geographical distribution of heterozygotes for mixed haplotypes of the most common African H2i and the predominant Eurasian H6i and H5j is quite revealing.

The population frequencies of the H2i/H6i genotype are widespread, although at low frequencies, not only in African populations, where its occurrence is 2-5%, but also in some Eurasian populations, where the frequency of this genotype is 1-2%.

Heterozygous carriers of the H2i/H5j haplotypes are more common in Africa than heterozygotes for H2i/H6i, the range of population frequency for H2i/H5j is from 5-7%, and in the Mandinka population of Gambia this genotype is the most common.

Carriers of the H2i/H6i genotype have also been found in all populations of East Asia, Western Europe and some European populations of Northern Eurasia with a population frequency of 1-2%.

The H2i/H4i genotype is found in African populations with a frequency of 3-5%, with only the Mandinka having a frequency of about 2%. This genotype was also found in the southern East Asia and in the British and Italian populations of Europe with a frequency of 1-2%.

### Population characteristics of ATM haplotypes in Africa

Haplotypes H1j, H1i, H2j, H3i, H4j, and H5i are exclusively characteristic of the African continent. H4i is third in frequency in Africa (from 9% among Yoruba to 13.5% among Mandinka and Mende) but has been detected outside Africa only in the Bashkir population at a frequency of 1%. This observation likely reflects the heterogeneity of this population, which was formed through the contribution of various ethnic groups.

The haplotype H1j occurs in African populations with frequencies ranging from 3% to 9% (on the western coast) and differs from the ‘ancestral’ one only at a single position (G in rs645485). H1j was found in the homozygous state exclusively among the Mende from Sierra Leone, where the frequency of homozygous carriers reached ∼3.5%. H1i is most prevalent on both the eastern and western African coasts (7-8%) and almost completely absent in the central part of the continent (0-1%). Homozygous carriers of H1i were detected with a population frequency of ∼1% among Mandinka from Gambia and Luhya from Kenya.

H2j was found at a frequency of 1% in Luhya and Mende populations. H3i is rarest in the Mende population (∼1%), while its population frequency increases to 5-6% in Luhya and Mandinka populations on the eastern and western coasts.

H4j is most prevalent among Mandinka (∼7%) in western Africa, with a decreased presence of ∼1.5% observed in Luhya populations in the east. In contrast, H5i reaches its highest frequency on the eastern coast (∼10%), while in other African populations, its population frequencies range from ∼4-5%.

Overall, it can be concluded that the African continent exhibits maximum diversity among ATM haplotypes and unique patterns of their population distribution.

## DISCUSSION

The study of *ATM* haplotypes in human populations of Eurasia and Africa from the standpoint of the “yin-yang” concept showed the presence in all populations of a conservative LD-block consisting of 28 extra-exon loci of the *ATM* gene located throughout the DNA sequence of the gene, including 5’ and 3’ untranslated regions of the *ATM* gene. Analysis of the phylogenetic relationships between 11 identified *ATM* haplotypes revealed the presence of two independent branches of *ATM* gene evolution from the ancestral haplotype, leading to the formation and widespread distribution of haplotypes with mutually exclusive alleles according to the “yin-yang” principle. In the samples analyzed outside the African continent, the dominant haplotypes in all human populations were the H6i haplotype (from 21 to 56%) and the H5j haplotype (from 32 to 72%), while the population frequency of the remaining haplotypes did not exceed 7% in Eurasian populations. In the sample from the African continent, 11 haplotypes were found, reflecting all stages of gene evolution from the “ancestral” haplotype. The most common was the H2i haplotype, its median frequency in African populations was 23%. On the territory of Eurasia, the H2i haplotype was found in only 7 populations, where its occurrence did not exceed 2%, which may be the result of evolutionary and demographic processes. Combining the obtained information on the evolutionary variability of *ATM* haplotypes with data on their frequencies in modern populations of Africa and Eurasia, it can be assumed that the process of formation of existing *ATM* haplotypes occurred on the African continent. Phylogenetic analysis of *ATM* haplotypes demonstrates the existence of two isolated ancestral human populations in the past, followed by their merging into a single mixed population on the African continent (Figure 3). Similar conclusions were reached in the project [11], which was conducted using data from the human genome of the 1000 Genomes Project, as well as the genomes of Denisovans and Neanderthals - Eastern and Western. Dutta et al. found an abundance of mutually exclusive pairs of common haplotypes for population-frequent (>25%) polymorphic markers in the human genome. Moreover, these “yin” and “yang” haplotypes had close population frequencies in Eurasia. At the same time, this study identified a category of haplotypes that are mostly observed in Africa. These haplotypes are composed of fragments of “yin” and “yang” haplotypes, hence the term “mosaic” used by the authors.

The results of the Tajima test in our study confirm the fact of the probable division of the original human population into several independent groups, and the statistically insignificant values of D when calculating the Tajima test for the haplotypes of the “yin” and “yang” branches suggest a neutral nature of evolution for each of the branches (Table S1).

The combined mixed population of carriers of *ATM* haplotypes, based on phylogenetic data, continued to exist on the African continent. Only a part of the *ATM* haplotypes from the “African” set were found on the territory of Eurasia. As a result of genetic drift and, possibly, balancing selection, the “yin” and “yang” haplotypes became dominant in Eurasia, while the others either disappeared completely or survived in small quantities in individual populations. The peoples of Asia differ from those of European origin, firstly, by the widespread occurrence of the H3j haplotype (up to 7%) and, secondly, by the prevalence of the H6i haplotype in East Asia. It is important to note that in South Asia, the dominant haplotype among peoples of European origin, H5j, prevails, which allows us to consider the modal haplotype H5j in the population as a distinctive Indo-European feature. An analysis of literary data on the connection of the analyzed polymorphic loci with oncological diseases revealed one important contradiction. In populations of East Asian origin, the H5j haplotype, containing the rs189037, rs609261 and rs4585 loci, is associated with the manifestation of negative effects in oncopathology. In peoples of European origin, this association is not observed, which may be due to the presence of an additional factor, yet unknown, which acts as an additive in East Asian populations. It is also worth paying attention to the prevalence of heterozygotes for the “yin” and “yang” haplotypes in most Eurasian populations. There are scientific data on the positive effect of heterozygous carrier references between the GA genotype and longevity, which the authors associate with a moderate level of *ATM* protein expression [14]. A study by Ding et al. showed that heterozygous genotype for rs189037 is associated with a reduced risk of type 2 diabetes in older adults [15].

## CONCLUSION

Our study of evolutionary changes in the *ATM* gene haplotypes, represented by high-frequency polymorphic loci throughout the DNA sequence of this gene, indicates a very ancient, significant in time African period of isolation of two branches of the ancestral human population and the subsequent merging of these branches into a single population space on the African continent. Repeated migration events to the territory of Eurasia probably occurred after the unification of ancient isolated subpopulations. Some differences in the composition of *ATM* haplotypes in the east and west of the Eurasian continent, as well as in individual populations, are probably due to the effect of gene drift.

## ACKNOWLEDGMENTS

Authors express the gratitude to the “Biobank of North Eurasia” for providing DNA samples, and to E.V. Balanovskaya for her comments and suggestions that help improve the article. The research was carried out within the state assignment of MSHE of RF for RCMG.

## AUTHOR CONTRIBUTIONS

Marina V. Olkova – article idea, bioinformatics analysis, primary genogeographic analysis, writing the article text

Sergey M. Koshel – genogeographic analysis

Georgiy Yu. Ponomarev – statistical and cartographic analysis

Andrei A. Alimov – research supervision, article editing

## DATA AND CODE AVAILABILITY

The 1000 Genomes data can be obtained from the 1000 Genomes Project (Ensembl) Phase 3 data. The published article includes all data on Northern Eurasian populations analyzed during this study. R code is available at https://github.com/molkova/ATM-haplotypes/tree/main.

## FIGURES

**Figure S1.**
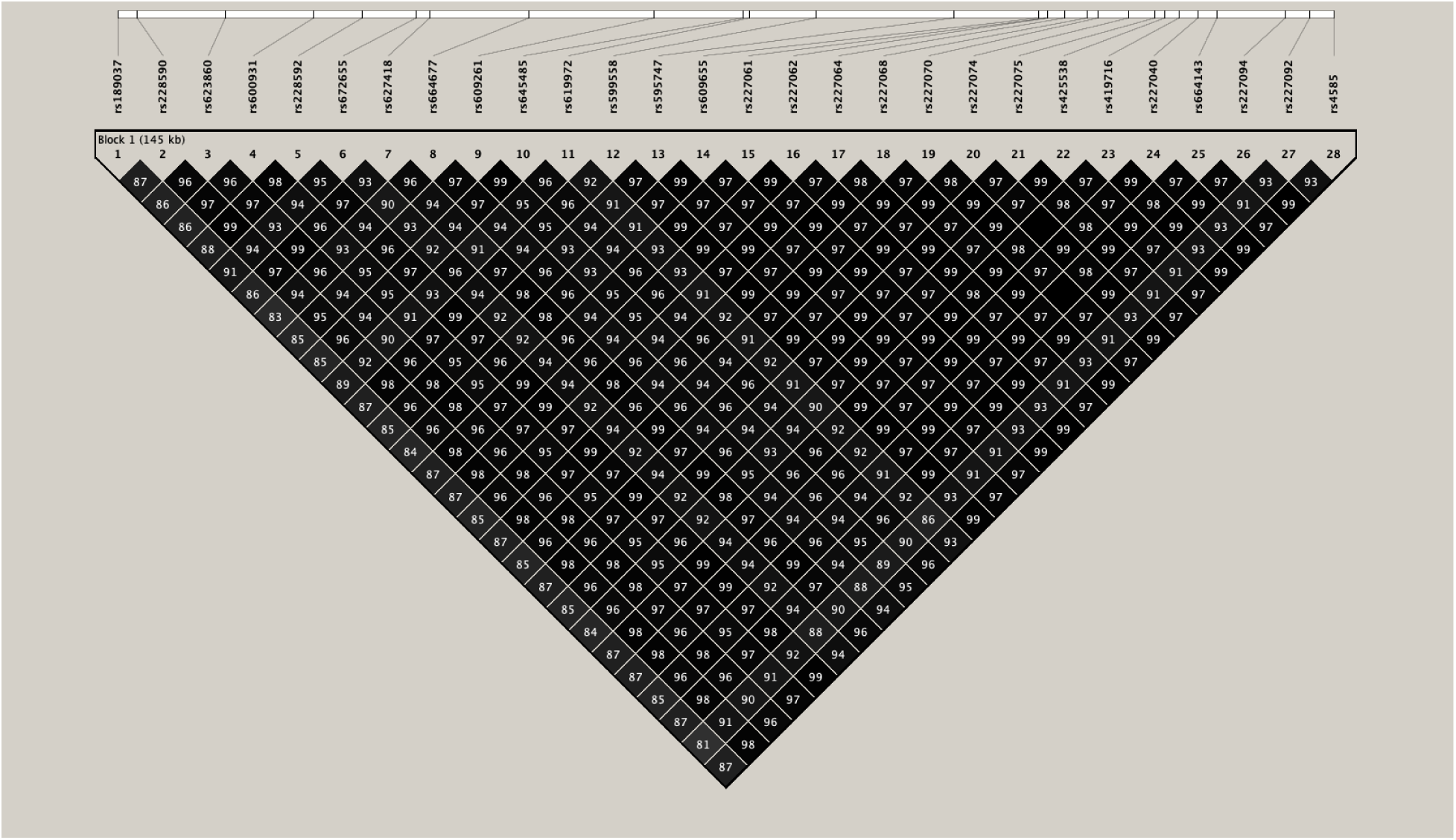
- LD-block of *ATM* SNPs. The numbers in the cells are the r^2^ value multiplied by 100.

**Figure S2.**
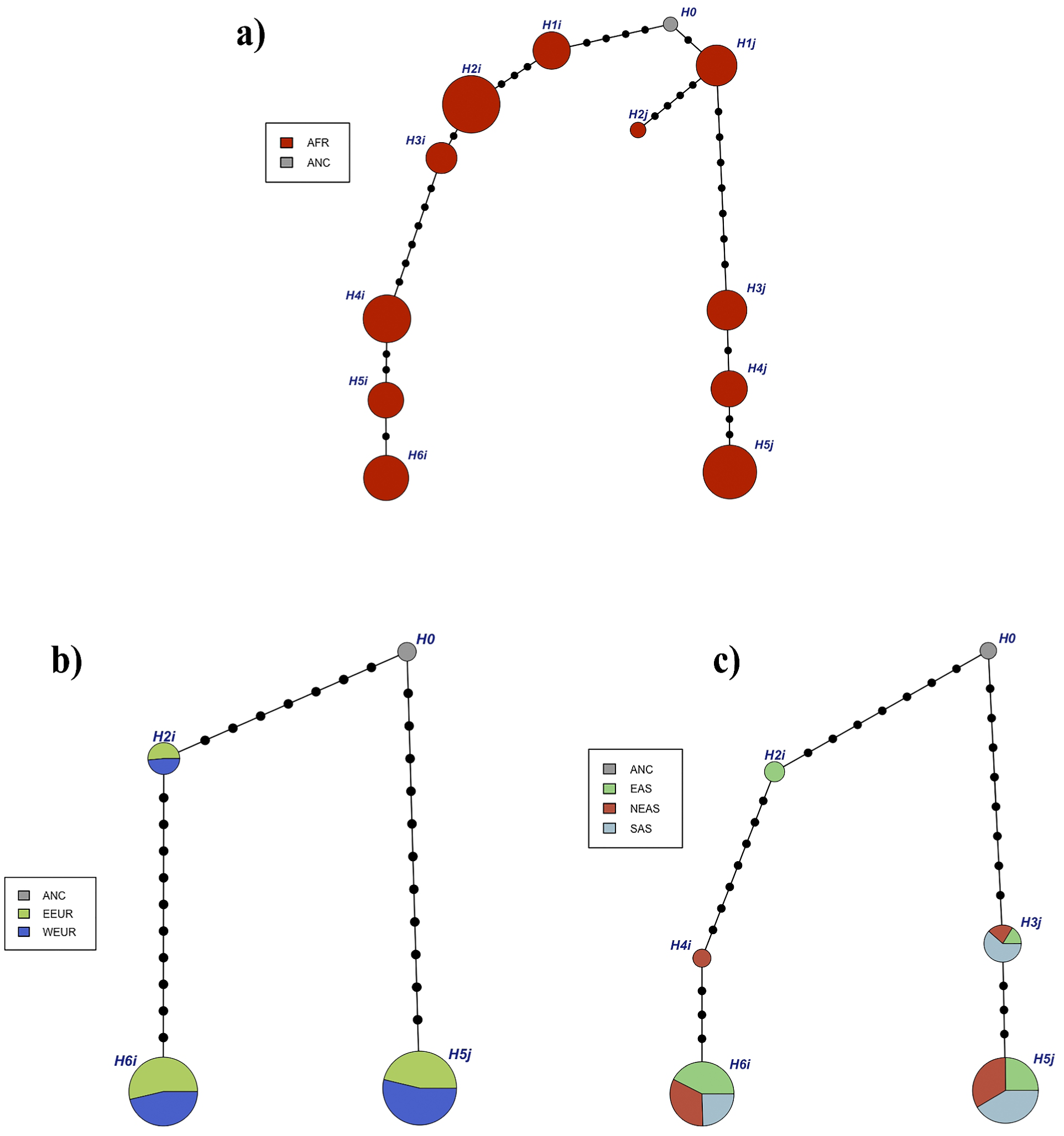
- Phylogenetic networks based on medians of population frequencies of *ATM* haplotypes for populations of African (a), European (b) and Asian (c) origin

**Figure S3.**
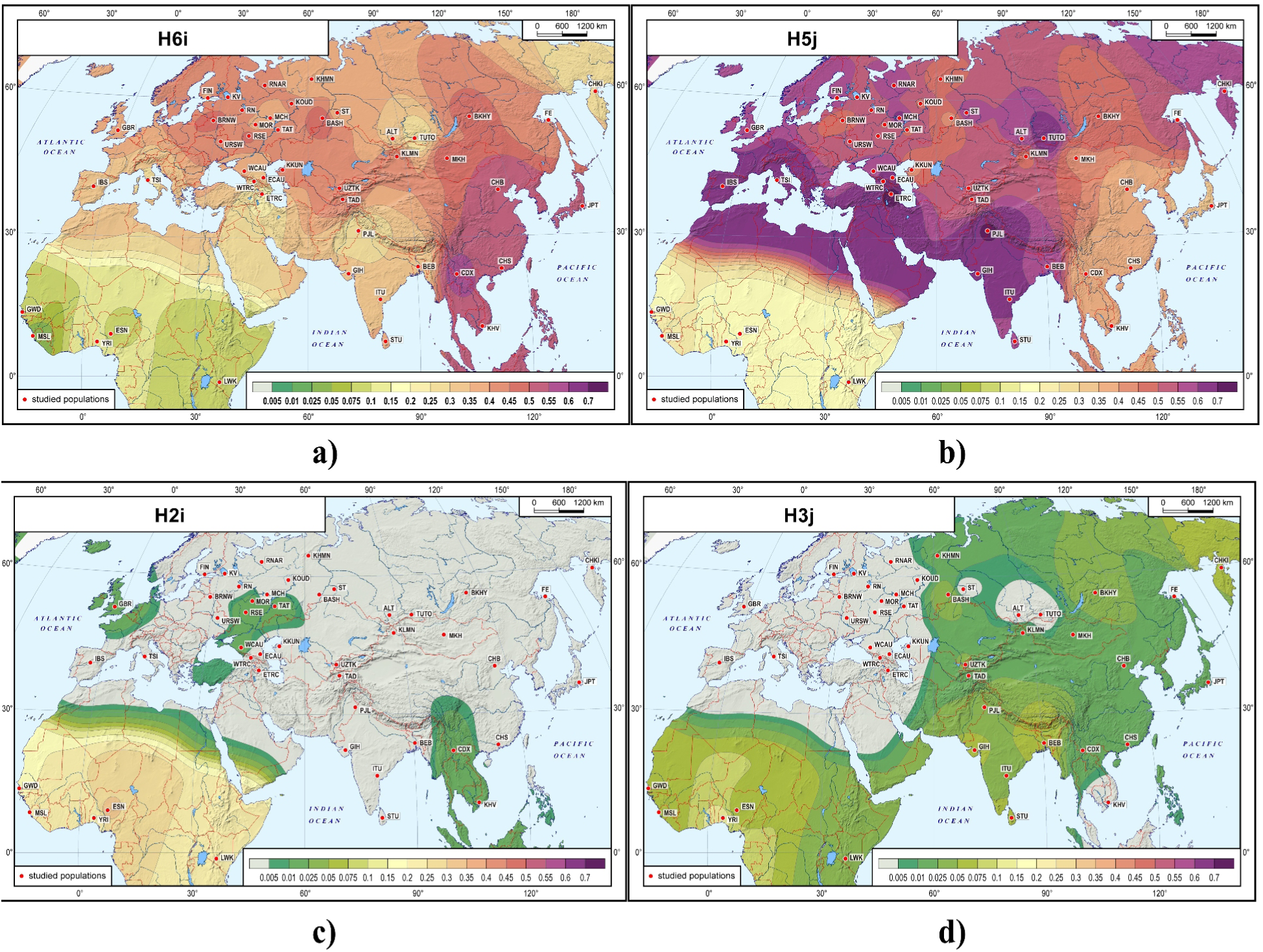
- Geographic distribution of the most common *ATM* haplotypes

**Figure S4.**
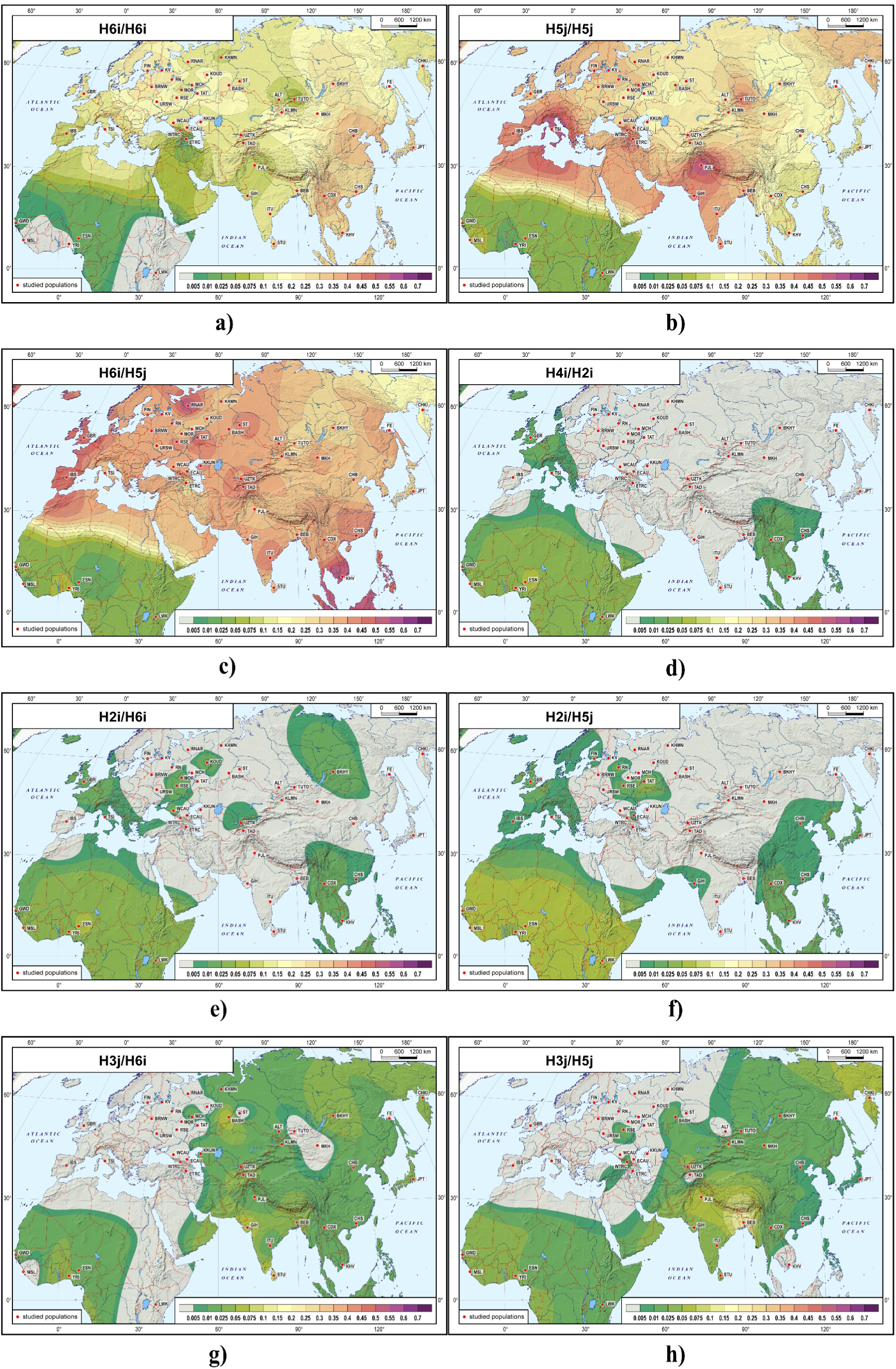
- Geographic distribution of the most widespread Eurasian *ATM* genotypes

**Figure S5.**
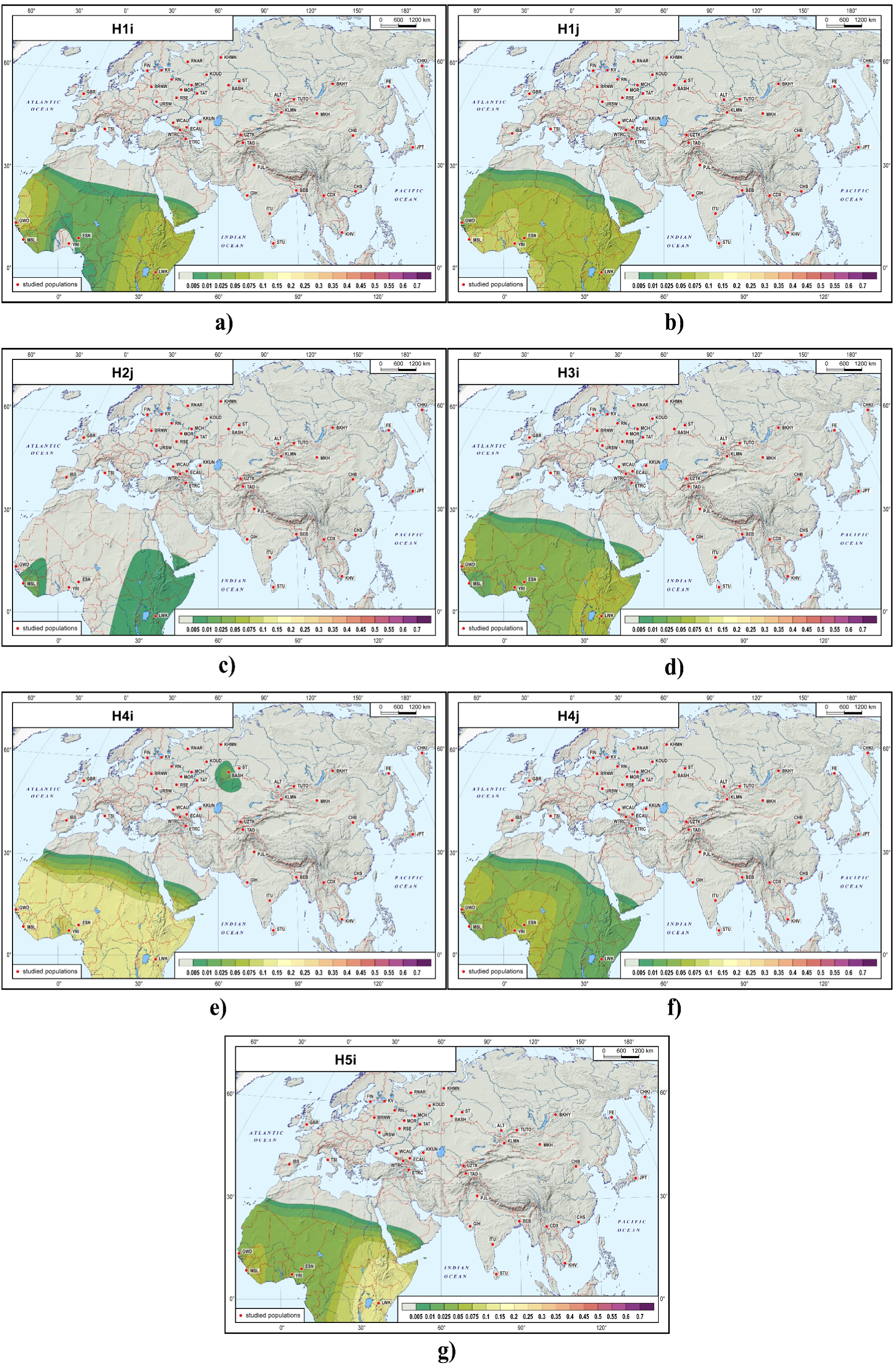
- Geographic distribution of African-specific *ATM* haplotypes

**Figure S6.**
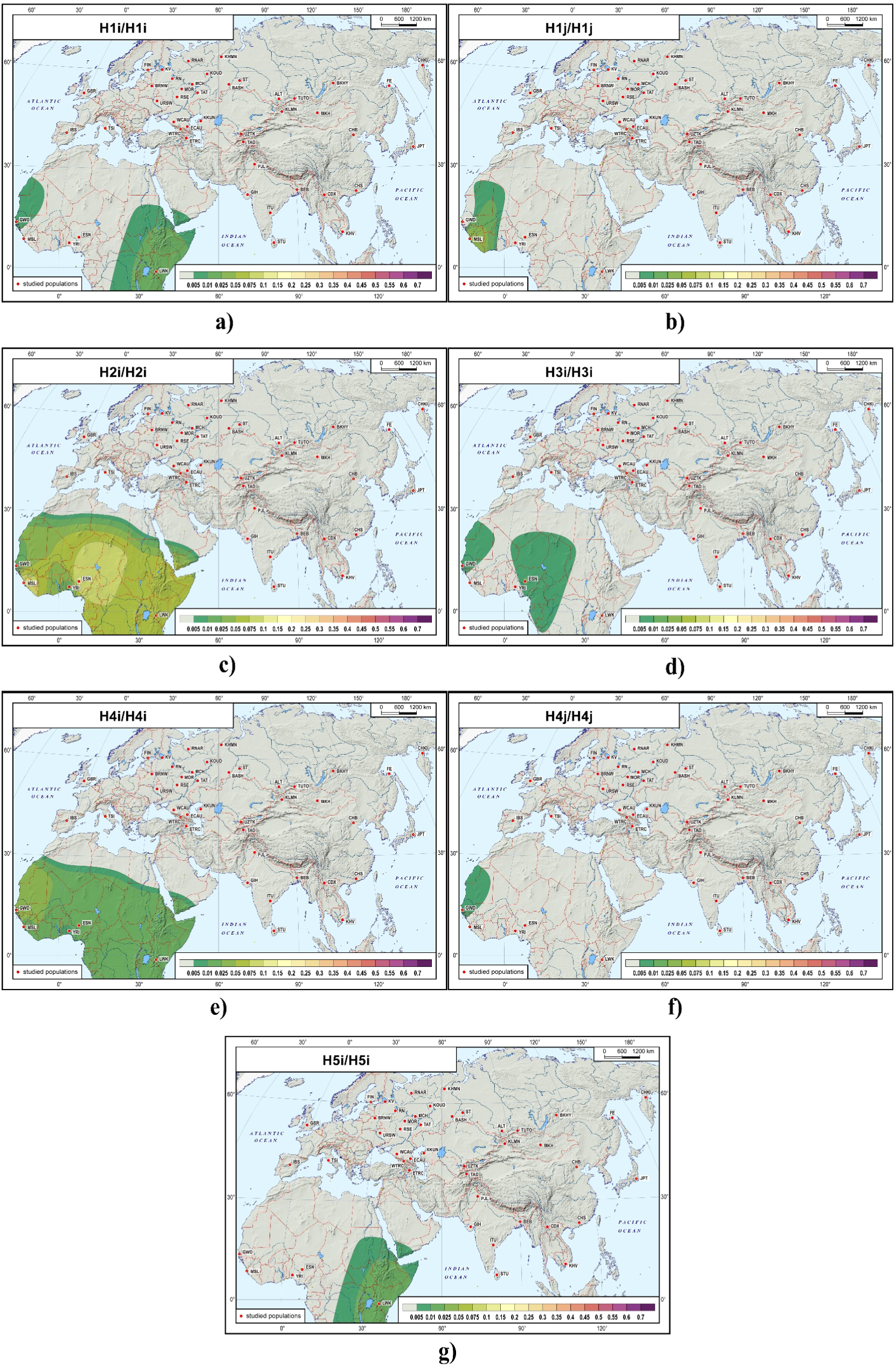
- Geographic distribution of *ATM* genotypes defined by African-specific haplotypes

## TABLES

**Table S1.**
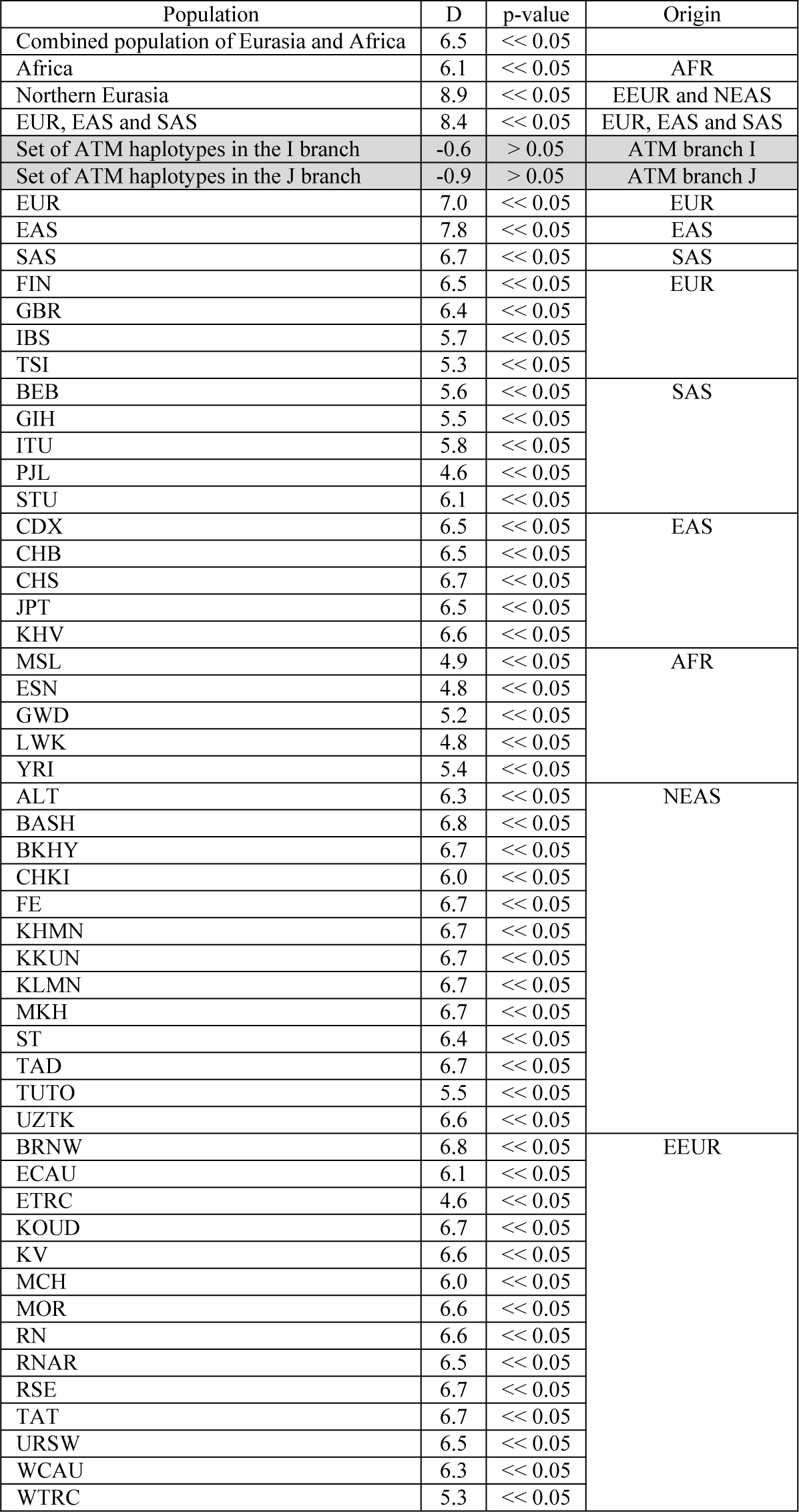
- Tajima’s test for *ATM* haplotypes. Abbreviations for populations can be found in Table S1. and for populations by origin: AFR – African, EAS – East Asian, EEUR – Eastern European, NEAS – Asian of Northern Eurasia, SAS – South Asian, WEUR – Western European

